# Evidence for Membrane Complex Assembly in Nanoelectrospray Generated Lipid Bilayers

**DOI:** 10.1101/661231

**Authors:** Matthias Wilm

## Abstract

Nanoelectrospray can be used to generate a layered structure consisting of bipolar lipids, detergent-solubilized membrane proteins, and glycerol that self-assembles upon detergent extraction into one extended layer of a protein containing membrane. This manuscript presents the first evidence that this method might allow membrane protein complexes to assemble in this process.

## 2. Introduction

Traditionally, electrospray is a tool to generate thin films of solid material^1^. It is possible to compose such films on the molecular level using nanoelectrospray^2^. This technique allows the generation of a layered structure that assembles into an extended sheet of a membrane containing outer membrane porin G (ompG) upon extraction of the ompG stabilizing detergent. Such films can be of interest as a starting point to manufacture biosensors and other technological devices that mimic properties of biological membranes. Often, a particular function in biological pathways is carried by a protein complex and not by a specific protein. Here we show the first evidence that within a nanoelectrospray generated layered structure proteins can assemble into complete membrane based complexes. When added to a nanoelectrospray preparation, listeriolysin O and pneumolysin form rings that insert themselves into lipid bilayers and form pores.

## 3. Materials & Methods

### 3.1. Protein Expression & Purification

Listeriolysin O (LLO) came from two sources. LLO was expressed and purified as described in^3^. Briefly, for expression in E.Coli BL21 (DE3) the gene segment encoding residues 25 - 529, the LLO sequence without the N-terminal secretion signal, complemented by an N-terminal His_6_ tag was cloned into the plasmid pET15b. The cells were grown overnight and then transferred into a selective medium containing 100 µg/ml ampicillin. After reaching a cellular density A_600_ of 1.2, the reduction of the temperature from 37°C to 30°C reduced the proliferation of the cells. The addition of 1 mM isopropyl β-D-1-thiogalactopyranoside inhibited the plasmid’s Lac-repressor and the protein expression starts. After 4 hours, the cells were collected, suspended in 50 mM Tris pH 7.7, 150 mM NaCl, and disrupted using a microfluidizer (M-110L, Microfluidics Corp., MA). After 1 hour centrifugation at 12 000 g, the supernatant was loaded onto a Ni-column to extract the His_6_-tagged protein. The protein was cleaved from the column with thrombin, concentrated on a 30 kDa cut-off membrane and purified with a gel filtration column (Superdex200) using 25 mM Tris (pH 7.7), 150 mM NaCl as running buffer. Recombinant LLO from Abcam (Cambridge, UK) was an alternative source for the protein (residues 60 - 529, product code ab83345).

The method to obtain pneumolysin (PLY) was similar^4^. An Escherichia Coli colony expressed the His_6_ tagged version of the protein by cloning it into the plasmid pET15b. The cell culture grew overnight at 37°C. Transfer to a selective medium containing 50 µg/ml ampicillin ensured that only plasmid containing cells survived. With reaching an A_600_ between 1 and 1.5, the temperature is reduced to 30° to reduce cell proliferation. The addition of 0.5 mM isopropyl β-D-1-thiogalactopyranoside induced protein expression. After overnight expression, the cells were harvested, suspended in lysis buffer and disrupted using a microfluidizer. It followed the centrifugation of the cells for 1 hour at 185 000 g and loading of the supernatant onto a HisTrap FF column. With a lysis buffer containing 500 mM imidazole to outcompete the histidine-tag from its Ni^+^ binding sites, the protein elutes from the column. Using a 50 mM Tris pH 7.0, 5 mM β-mercaptoethanol solution, the protein fractions are diluted. Overnight incubation with 100 unit Thrombin at 4°C removed the His tag. Passing the protein solution over an HiTrap Q FF using a salt gradient from 0 to 1 M NaCl in a 50 mM Tris, pH 7.0, 5 mM β - mercaptoethanol buffer resulted in a purified sample. The protein solution was not frozen but kept at 4°C to avoid precipitation.

### 3.2. Electron Microscopy

#### 3.2.1. Molecular Layer Inspection

Exposing carbon coated grids briefly to gas discharge renders them hydrophilic and ready to take up lipid layers covered by glycerol from an aqueous buffer solution. The transfer of the lipid films to the grids succeeds by touching the liquid meniscus briefly from above. Subsequently, several exchanges of buffer or doubly distilled water wash off the glycerol. A droplet of 1% wt/vol uranyl acetate is placed onto the target to increase the contrast for microscopic inspection. Alternatively, a platinum/carbon beam shot at an angle of 15° generates an up to 1.5 nm thick metallic layer on the surface of the grid. An FEI Tecnai Spirit BioTWIN transmission electron microscope operating at 120 kV with magnifications of up to 150 000x generated all the images.

### 3.3. Nanoelectrospray based Surface Preparation

#### 3.3.1. Electrospray Apparatus

A self-built electrospray apparatus was used to spray molecular layers onto a liquid buffer meniscus with a diameter of 3 mm. A gold coated glass capillary with one end pulled to a sharp tip with a glass capillary puller (model P-87 Puller, Sutter Instruments Company, Novato, CA, USA) contains the liquid sample to be electrosprayed^2,5^. A gas-tight holder fixes the capillary. A small metal plate whose distance to the emitter can be changed centers the buffer-containing cylinder under the spray. A voltage of up to 10 kV initiates the electrospray after breaking the tip of the capillary to an orifice size of about 1 µm^6,7^. The distance between the electrospray emitter and the buffer surface is in the order of 3 cm. This flight path is long enough for the solvent from the electrospray droplets to evaporate so that only non-volatile components reach the buffer-meniscus. The flow rate is not directly controlled but varies between approximately 20 nanoliters/min and several hundred nanoliters/min depending on the size of the orifice and the applied pressure.

#### 3.3.2. Layer Preparation

The buffer for the pneumolysin and listeriolysin O preparation was a 50 mM Tris, 150 mM NaCl aqueous solution. For the listeriolysin O solution, the pH was adjusted to 5.5, for of the pneumolysin solution to 7.0. The first step in generating protein filled membrane layers is to spray 1 *µ*l of 66 pmol/*µ*l lipid solution with 35% cholesterol and 65% phosphatidylcholine in ethanol onto the meniscus of the buffer. Overnight incubation finalizes the formation of a closed lipid bilayer. A thin glycerol layer generated by spraying a 1:2 solution of glycerol/ethanol for approximately 10 minutes creates a hydrophilic base for the subsequent membrane assembly.

The next step consists of adding either a half or a complete lipid bilayer. As before, incubation overnight completes bilayer formation. Now, the surface is prepared to add the protein containing film. The pneumolysin solution consisted of 26 pmol/*µ*l protein in 50 mM Tris, 150 mM NaCl at neutral pH and the listeriolysin O solution had a concentration of 20 pmol/µl dissolved in 50 mM Tris, 150 mM NaCl at ph 5.5. The solution sprayed had an additional component of 25% glycerol. When the surface carried only a lipid monolayer, the protein solution contributes enough lipid to complete the bilayer - 20 pmol/*µ*l lipid, 35% cholesterol, and 65% phosphatidylcholine. About 1 *µ*l of the final solution distributed over the 3 mm large liquid meniscus generates the protein containing layer. Spraying a 1:2 glycerol/ethanol solution for 10 minutes finalizes the hydrophilic environment. The self-assembly process proceeds throughout the incubation period of several days at room temperature.

## 4. Results

Listeriolysin O and pneumolysin are both members of a group of pore-forming toxins. They are soluble 53 kDa respective 60 kDa proteins. Up to 50 monomers form a non-covalent, ring-shaped complex on membranes, insert themselves, and create large pores of about 30 nm in diameter^3,8–10^. Listeriolysin O assembles best under the slightly acidic pH of 5.5. Both proteins require phospholipids in the membrane for docking and complex formation.

### 4.1. Listeriolysin O Assembly on a Lipid Bilayer

The buffer chosen for the preparation of listeriolysin O layers allows the assembly of its complexes on membranes^11^. However, in these experiments, the local environment for the protein is a thin glycerol layer and not the buffer itself. In thin layers, ion concentrations and pH can be very different from the bulk solution. In an experiment to test the complex formation, the nanoelectrospray generated first a lipid bilayer film, then a listeriolysin O containing film followed by a glycerol layer to finalize the hydrophilic environment. **Figure 1** shows the result. The overall appearance corresponds to published listeriolysin O complexes on membranes^11^. Remarkable are the electron-dense centers of many of the completed rings. They might be attributable to remnants of glycerol and salt residues from the buffer solution that the washing procedure did not successfully remove. Mono-molecular ompG pores in a nanoelectrospray prepared layer have a similar appearance^2^.

**Figure 1.**
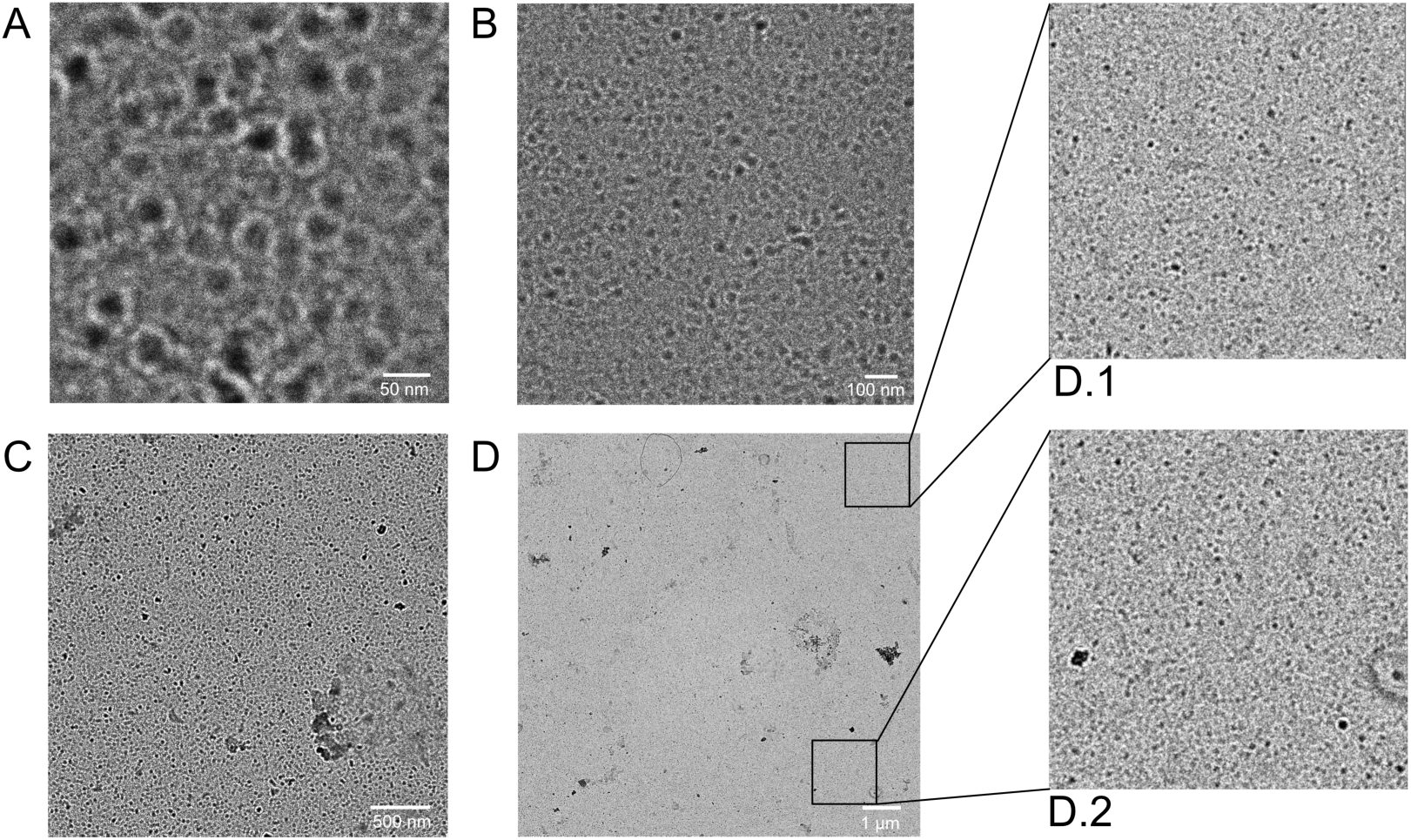
Transmission electron microscopic image of listeriolysin O sprayed onto a lipid bilayer and covered with glycerol: A negative stain with 1% uranyl acetate solution increases the contrast. A large layer of listeriolysin O assembly is visible. The white bar on panel A is 50 nm long, on panel B 100 nm, panel C 500 nm and on panel D 1000 nm. Pores assembled in supported lipid bilayers have a comparable density^12^.

### 4.2. Listeriolysin O and Pneumolysin Assembly on a Lipid Monolayer

The physiological mechanism of listeriolysin O and pneumolysin is to assemble to a ring-shaped complex onto a bilayer membrane, to insert and to form a pore^4,11^. In the context of nanoelectrospray generated layers, however, it is of interest whether the layered structure allows the formation of protein complexes of intra-membranous proteins. An ompG containing membrane only assembles when the basis for the protein layer is a lipid monolayer^2^. **Figure 2** shows the result of an experiment when spraying pneumolysin onto a lipid monolayer. The images show a strand of precipitated salt from the buffer with pneumolysin. Pneumolysin dissolves into the surface layer, diffuses away and, step by step assembles into a pore structure. A pore formation in a supported bilayer has been directly observed using atomic force microscopy for the membrane attack complex (MAC)^13^. The pore completes in about 100 seconds. Here, in this experiment, the process probably takes more time since diffusion in a thin glycerol layer is slower than in an aqueous solution.

**Figure 2.**
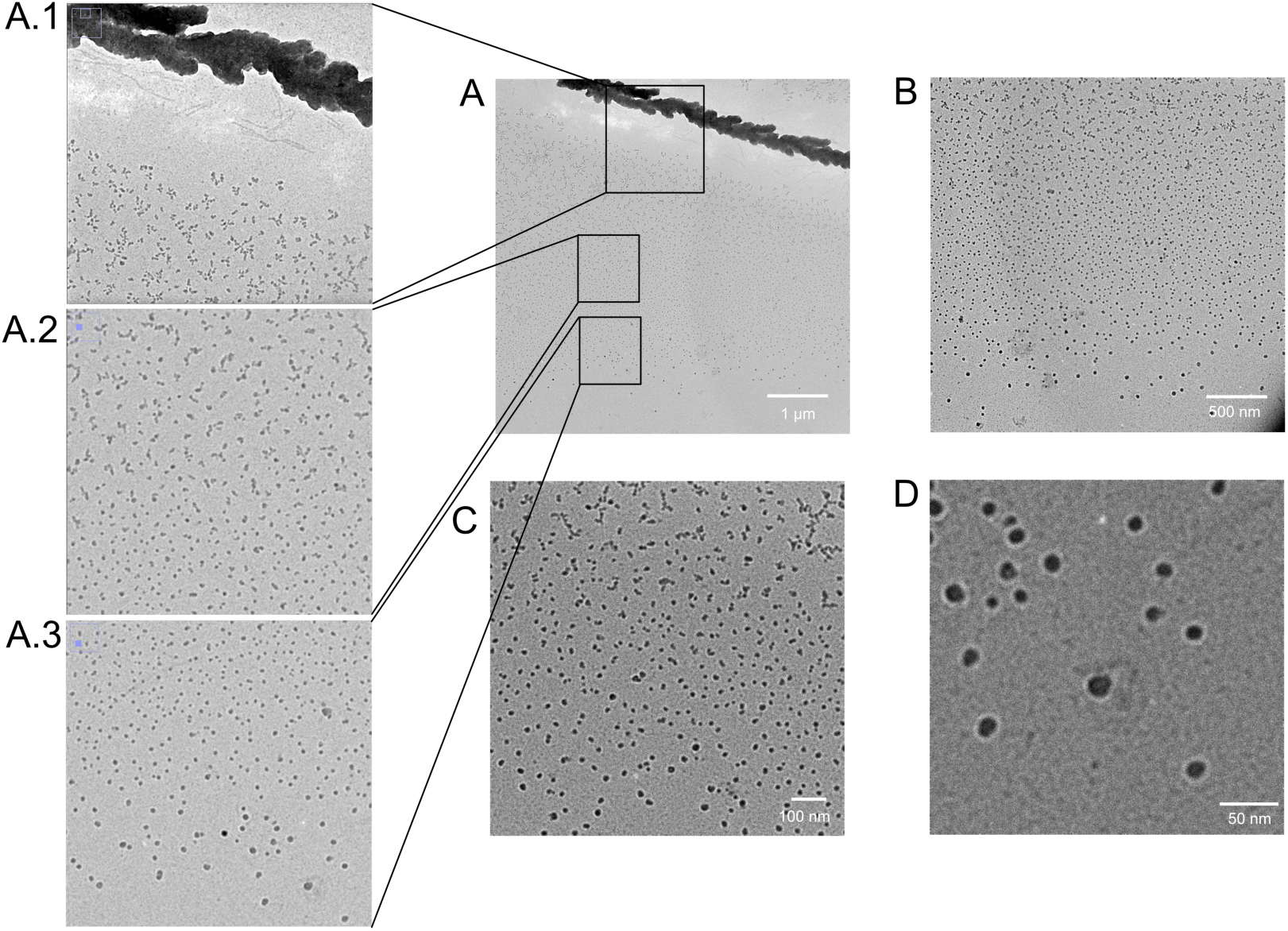
Pneumolysin preparation on a lipid monolayer: pneumolysin was sprayed onto a lipid monolayer and covered with glycerol. After five days incubation, the layer was transferred to a carbon coated grid, washed with buffer, negatively stained with 1% uranyl-acetate and washed with water. The scale bar in panel A is 1 µm long, in panel B 500 nm, in panel C 100 nm, and in panel D 50 nm.

The pneumolysin pores displayed in **Figure 2** have the expected size^4^ but show an electron-dense center similar to ompG prepared in a similar way^2^. A platinum-shadowed preparation should yield a clearer picture of the observed structure. **Figure 3** shows a typical result. The first observation is that, corresponding to the mechanism for listeriolysin O and pneumolysin pore formation, the efficiency of pore assembly when providing only a lipid monolayer as a base is much lower than when spraying them onto a lipid bilayer as shown in **Figure 2**. When using either pneumolysin or listeriolysin O, the round structures appear in the same way. They are flat since they don’t have any shadow in a platinum-sputtered preparation. They are absent from control experiments when omitting the protein. Moreover, using listeriolysin O from a completely different source, purchased from Abcam, the same structures appear.

**Figure 3.**
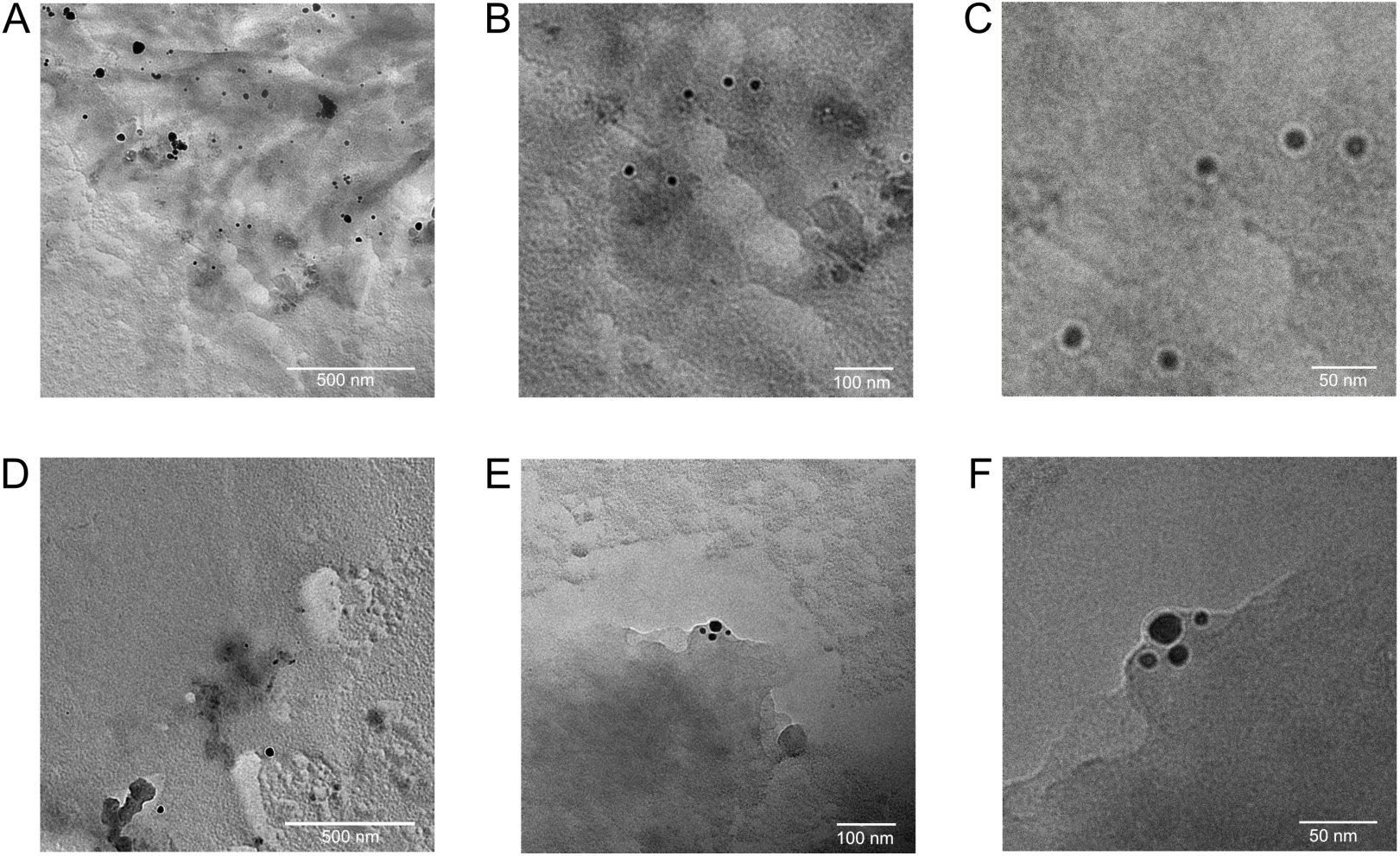
Pneumolysin and Listeriolysin O sprayed onto a lipid monolayer: panels A, B, and C show a pneumolysin preparation, panel D, E, and F a listeriolysin O experiment. The grids were not stained but shadowed with a 1.5 nm thick platinum layer at an angle of 15°. They do not protrude from the surface since they do not have a visible shadow. When compared with the on-membrane structures as displayed in **Figure 1**, they are not numerous. The scale bars of panel A and D are 500 nm long, panel B and E 100 nm and panel C, F 50 nm.

## 5. Conclusions

Listeriolysin O, sprayed onto an intact lipid bilayer and embedded into glycerol, can dock onto the membrane and assemble into the classical pores (see **Figure 1**). This observation adds evidence that a bilayer forms when the nanoelectrospray distributes the appropriate amount of lipids onto a liquid meniscus. The protein maintains it’s basic structure throughout the preparation within a flat glycerol layer. Exposing the protein to fewer lipid molecules, to the equivalent amount of a single monolayer, creates an unphysiological situation and leads to different results. The purpose of this experiment was to gather at least indirect evidence whether nanoelectrospray prepared layers will allow membrane protein-based complexes to form. OmpG as an integral membrane protein integrated only under the condition that it had a lipid monolayer as base^2^.

**Figure 2** shows the first result obtained. The images seem to display the assembly process of pneumolysin to pore-like structures. The pores have the same size as physiological pneumolysin pores but look different than those assembled into lipid bilayers^4^. However, the assembly imaged in **Figure 2** does not occur on lipid bilayers and, therefore, does not reflect a physiological situation. Similar to ompG pores they have an electron-dense center that might be caused by not washing out the glycerol entirely^2^. Water cannot easily penetrate holes with sizes in the nm range when washing the surface layer on an electron microscopy grid.

To further investigate the nature of these structures assembled from pneumolysin, additional preparations were made to visualize them after platinum shadowing (see **Figure 3**). Experiments conducted with pneumolysin and listeriolysin O from two different sources confirmed the observation. The rings have the same size as listeriolysin and pneumolysin pores in bilayers. The electron-dense centers do not surpass the surface similar to ompG based preparations. However, in contrast to assemblies observed on bilayers (see **Figure 1**), they are not numerous.

In summary, the results displayed in **Figure 1** and **Figure 2** seem to show that nanoelectrospray based layer preparation of membranes and proteins can allow the assembly of membrane-protein complexes. The evidence can only be indirect since listeriolysin O and pneumolysin are soluble proteins. However, accepting this limitation, the demonstrated results encourage further experiments with more complex membrane-based protein structures.

## 7. Acknowledgments

All experiments were done in the laboratory of Prof. W. Kühlbrandt at the Max Planck Institute of Biophysics, Frankfurt, Germany. I am grateful to Prof. Kühlbrandt for providing the necessary support to conduct all these experiments. I thank Dr. Oezkan Yildiz and Dr. Katharina von Pee for the intellectual and material assistance. Without the help of Deryck Mills, the project could never have been done.

The work of Matthias Wilm was supported by the Science Foundation Ireland, SFI grant 07/SK/B1184c.

